# Accurate *ΔT_m_* Prediction Without Protein Structure Inputs for Biomolecular Stability

**DOI:** 10.64898/2026.07.02.735991

**Authors:** Daniel Siegismund, Mario Wieser, Eriberto Natali, Stephan Steigele

## Abstract

Predicting protein stability, like changes in melting temperature (*ΔT*_*m*_) caused by mutations, is a critical task in therapeutic protein engineering and drug discovery. This is reflected by a growing solution space, including both AI-based sequence and structure based methods. This paper demonstrates that accurate *ΔT*_*m*_ prediction does *not* require structural input features, but can achieve state-of-the-art results with a careful training design for large sequence-based protein language models. We combine an autoresearch-inspired setup search with controlled ablation studies and show that a well-tuned sequence-only ESM2-650M model [6] outperforms structure-informed methods in our benchmark, achieving the lowest error (MAE/RMSE) and competitive Pearson correlation without pH or structural inputs. We further show that choices such as loss function, pooling strategy, auxiliary supervision, and finetuning regime materially affect performance.

## 1 Introduction

Thermal stability is a critical developability attribute for protein therapeutics, such as monoclonal antibodies. It contributes to the ability to retain structural integrity during manufacturing, storage, transport, and in vivo use. Poor thermal stability can affect quality, efficacy, and safety [4]. Engineering antibodies to optimize for this property is a challenging task, requiring multiple trial and error rounds, resuming in long engineering time and high costs [3]. AI-based prediction models trained on thermostability data can however aid engineering phases [3] and inform pre-selection decisions for mutant design prioritization [10]. In particular, variations in Melting Temperature, also defined as (*ΔT*_*m*_) predictions, can anticipate whether mutations may stabilize the structure or not. *ΔT*_*m*_ prediction is a distinct task from *ΔΔG* (change in folding free energy) prediction and presents its own challenges. Recent works have explored increasingly complex model architectures (often incorporating protein structural information or multiple inputs), but it remains unclear whether such complexity or structural features are necessary for accurate *ΔT*_*m*_ prediction. In fact, comparisons in the literature often entangle architectural choices with training protocols and tend to optimize for only a subset of metrics, making it difficult to discern which factors truly drive performance improvements. Traditional methods for *ΔT*_*m*_ prediction, such as HoTMuSiC [11] and AUTO-MUTE [8], relied on protein structure features or statistical potentials combined with machine learning. However, deep learning methods for *ΔT*_*m*_ have been relatively scarce, likely due to limited data. Language models of protein sequences have recently been shown to capture structural and functional information from sequences alone, as demonstrated by large-scale models like ESM-1b and ESM-2 [13, 6]. Recent evidence further suggests that such models may already encode much of the predictive signal needed for *ΔT*_*m*_ prediction, with structural inputs offering limited additional benefit under current integration strategies [9]. As such, sequence-based *ΔT*_*m*_ prediction offers a scalable and interpretable route to de-risk antibody leads and guide engineering decisions across the discovery pipeline.

In this work, we show that a carefully tuned training strategy and simple design choices can have a significant impact on *ΔT*_*m*_ prediction performance, even when using a sequence-only pre-trained model backbone. We demonstrate that a large protein language model (ESM2-650M) with no structural inputs can achieve state-of-the-art accuracy on a standard stability benchmark. Our contributions are threefold: (1) an autoresearch-inspired setup search procedure to identify a strong reference recipe for *ΔT*_*m*_ prediction, (2) validation-only ablation experiments around this reference configuration that isolate the effects of loss function, auxiliary supervision, and pooling strategy, and (3) a demonstration that full fine-tuning of the language model yields the best results, while partial fine-tuning of top layers offers a favorable trade-off between performance and efficiency.

## 2 Related Work

Early approaches to *ΔT*_*m*_ prediction relied on structure-based features and classical machine learning. Methods such as AUTO-MUTE [8] and HoTMuSiC [11] combine structural representations with statistical potentials or shallow models, achieving reasonable accuracy but requiring experimentally resolved protein structures and limited datasets.

Recent work has shifted toward deep learning models that leverage sequence and structural information. Approaches such as DPStab [15] or GeoDTm [17] use transformer-based architectures for end-to-end prediction, while other methods incorporate multimodal representations combining sequence embeddings and structural features [18]. These models often improve predictive performance but introduce significant architectural complexity and reliance on additional inputs. Protein language models (pLMs), including ESM-1b and ESM-2 [13, 6], enable sequence-only prediction by encoding structural and functional information directly from amino acid sequences. Transfer learning with pLM embeddings has been successfully applied to thermostability prediction tasks [12], offering scalability and avoiding dependence on structural data.

## 3 Method

### 3.1 Preliminaries

We build upon the Evolutionary Scale Modeling (ESM2) family of protein language models introduced by Lin et al. [6] and the work of [18]. ESM2 is a transformer-based masked language model trained on large-scale protein sequence corpora using a self-supervised objective. Given an amino-acid sequence *s* = (*s*_1_, …, *s*_*L*_) of length *L*, ESM2 produces contextualised representations *H* ∈ R^*L*×*D*^, where *D* denotes the embedding dimensionality. Each embedding *H*[*i*, :] captures both local biochemical context and long-range interactions. In addition to per-residue representations, ESM2 outputs a special cls token embedding that serves as a global sequence representation.

### 3.2 Model

Let *s*^wt^ denote a wild-type protein sequence and *s*^mut^ its corresponding single-point mutant, differing at position *j*. Both sequences are tokenised and processed jointly by a shared ESM2 encoder, yielding contextual representations

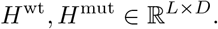

From these representations, we extract three types of features: (i) the global cls embeddings 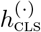, (ii) mutation-site embeddings 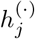, and (iii) mean-pooled sequence representations

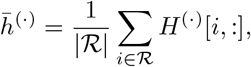

where ℛdenotes the set of valid (non-special and non-padding) residues.

Our predictor follows a two-head architecture that decomposes the prediction into complementary local and global components. The final prediction is obtained by averaging the outputs of the two heads.

#### Local Mutation Effect Head

The first head is designed to capture localised mutation effects by modelling second-order interactions between wild-type and mutant representations at the mutation site. Let 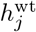, 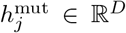 denote the corresponding embeddings. Direct computation of the outer product in the original space as proposed in [18] incurs a prohibitive *O*(*D*^2^) dimensionality. To address this, we introduce a *bottleneck projection* via learnable linear maps *W*_m_, 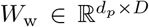with *d*_*p*_≪ *D* (here *d*_*p*_ = 128), yielding compressed representations prior to interaction:

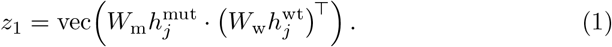

This projection reduces the feature dimensionality from *D*^2^ to 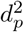, acting not only as a computational optimisation but also as an implicit regulariser that suppresses high-frequency noise in the pretrained representations. The resulting vector 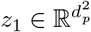 is processed by a neural network *f* :

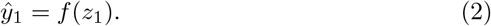

#### Global Difference Head

The second head is designed to capture global, sequence-level effects induced by mutation, and is explicitly decoupled from local substitution signals. While the formulation of Zhang et al. [18] combines mutation-site and global differences within a single feature space, we instead enforce *head specialisation* by restricting this head to operate exclusively on global representations. Concretely, we construct

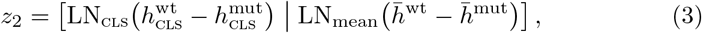

where LN denotes a layer normalisation that is applied independently to stabilise feature scales. By excluding mutation-site features, this head is encouraged to model distributed structural and thermodynamic effects, thereby eliminating redundant learning across the two heads and improving complementarity. The feature vector *z*_2_ is then passed through a neural network *g*:

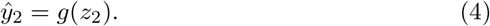

Finally, the loss is defined as:

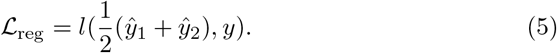

where *l* is the loss function, e.g. *l*_1_, and *y* is the ground truth.

#### Optional Auxiliary Classifier

A key challenge in *ΔT*_*m*_ prediction arises from the highly imbalanced target distribution, in which the majority of samples are concentrated. More specifically, there is a narrow range of mildly destabilising effects, while stabilising mutations and extreme destabilising cases occur relatively infrequently. As a result, the optimisation process is dominated by the high-density region of the target space, encouraging the model to prioritise accuracy in this regime while under-representing rare but informative cases. This leads to a bias towards central predictions and reduced sensitivity in the tails of the distribution. To mitigate this effect, we optionally introduce an auxiliary classification objective defined over discretised stability classes *c* ∈ {destabilising, neutral, stabilising}, constructed using a fixed threshold on *ΔT*_*m*_.

Let 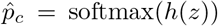 denote the predicted class probabilities produced by an auxiliary classifier *h* given the concatenated feature representation *z* = [*z*_1_, *z*_2_], and let *y*_*c*_ ∈ {0, 1}^3^ denote the one-hot encoded ground-truth class label. The auxiliary objective is optimised using the cross-entropy loss:

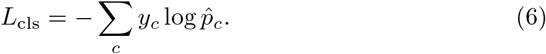

##### Algorithm 1

*ΔT*_*m*_ Prediction Algorithm

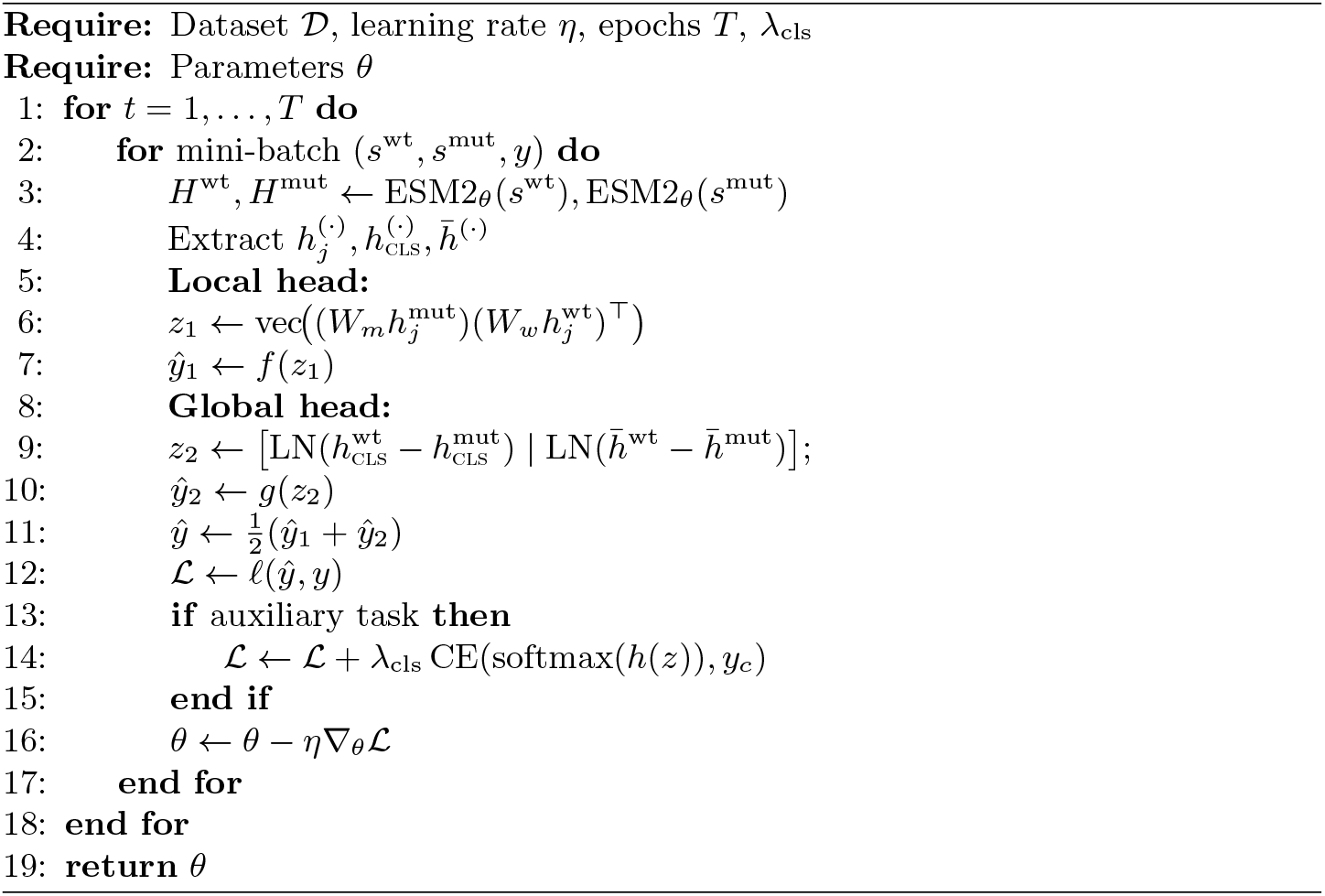

This auxiliary supervision encourages the model to better distinguish between coarse stability regimes, thereby improving sensitivity to rare stabilising and strongly destabilising mutations.

### 3.3 Objective function

The overall training objective is given by

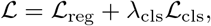

where *L*_cls_ denotes a cross-entropy loss and *λ*_cls_ controls the relative contribution of the auxiliary term. In contrast to Zhang et al., we employ robust alternatives to mean squared error (MSE), specifically *ℓ*_1_ or Huber loss. Unlike MSE, which disproportionately emphasises large residuals due to its quadratic scaling, these losses exhibit reduced sensitivity to outliers and heavy-tailed noise in the target distribution. This property is particularly desirable in the present setting, where extreme stability changes are rare but may correspond to noisy or heterogeneous measurements. Please refer to Algorithm 1 for a detailed description of the algorithm.

## 4 Experiments

### 4.1 Datasets

To enable direct comparison with prior work, we follow the data protocol of the GeoStab benchmark [17]. The training set (`S4346`) comprises 4,346 single-point mutations across 349 proteins, collected from ProThermDB and ThermoMutDB. The held-out test set (`S571`) contains 571 mutations across 37 proteins from the same sources.

Because the benchmark does not define an explicit validation split within `S4346`, we construct one for model selection following the reference protocol. Specifically, protein sequences are clustered with MMseqs2 [14] at 50% sequence identity, and clusters are partitioned into training and validation sets in an 8:2 ratio. All model selection, hyperparameter tuning, and design decisions are performed exclusively on this split. Final results are obtained by retraining on the combined training and validation data while preserving the original `S4346/S571` evaluation protocol.

### 4.2 Training Details

We adopt an *autoresearch-style* iterative procedure, inspired by [5], to identify a strong training configuration for *ΔT*_*m*_ prediction. This process operates entirely on the train/validation split derived from `S4346`, and the held-out test set `S571` is accessed only after the full training recipe is fixed. Starting from a standard ESM2-650M fine-tuning baseline, we iteratively refine the training setup by proposing and evaluating targeted modifications. The final training configuration employs AdamW [7] with *β*_1_=0.9, *β*_2_=0.98, and weight decay 0.03. Separate learning rates are applied to the backbone and prediction heads (1.5× 10^−5^ and 3 ×10^−4^, respectively), together with a OneCycleLR schedule comprising 30% linear warm-up followed by cosine annealing. We use dropout (*p*=0.2), gradient clipping (∥ **g** ∥ =0.1), batch size 8, and `bfloat16` mixed precision. Models are implemented in PyTorch and fine-tuned end-to-end unless otherwise noted. Unless stated otherwise, the final reported test model uses ESM2-650M with full finetuning, mean pooling, the two-head predictor, and *l*_1_ regression loss without auxiliary classification, as selected from validation performance.

### 4.3 Evaluation Metrics

We evaluate regression performance using Pearson correlation coefficient (PCC), mean absolute error (MAE), and root mean squared error (RMSE) between predicted and experimental | *ΔT*_*m*_|. To assess performance on challenging cases, we additionally report Tail MAE, defined as the MAE computed on the 10% of mutations with the largest *ΔT*_*m*_ in the validation set. All reported results are averaged over multiple random seeds and presented as mean ± standard deviation. After fixing the full training recipe, the model is retrained on the combined training and validation data for final evaluation on `S571`.

## 5 Results

We compare our method against prior approaches on the *ΔT*_*m*_ benchmark (Table 1). Our sequence-only model achieves the lowest MAE (4.964 ^°^C) and RMSE (7.430 ^°^C) among all methods, despite not using pH or structural information. While DPStab achieves the highest PCC (0.592), it requires pH input and yields higher MAE and RMSE. Overall, our results demonstrate that a sequence-only model can match or outperform structure-aware and multimodal approaches on this task.

**Table 1.** Comparison on the *ΔT*_*m*_ benchmark (test set). “Struct.” indicates whether the method uses explicit or predicted structural information.

| Method | PCC↑ | MAE↓ | RMSE↓ | pH | Struct. |
| --- | --- | --- | --- | --- | --- |
| HoTMuSiC [11] | 0.33 | 5.70 | 8.41 | No | Yes |
| AUTO-MUTE [8] | 0.29 | 5.79 | 8.50 | No | Yes |
| Geo$\Delta T_m$-Seq [17] | 0.46 | 5.55 | 8.11 | No | No |
| Geo$\Delta T_m$-3D [17] | 0.535 | 5.308 | 8.034 | Yes | Yes |
| ESM3-$\Delta T_m$ [18] | 0.50 | 5.21 | 7.68 | No | Yes |
| DPStab [15] | **0.592** | 5.050 | 7.643 | Yes | Pred. |
| **Ours** | 0.553 | **4.964** | **7.430** | No | No |

### 5.1 Ablation Studies

#### Backbone

We evaluate different protein language model backbones (Table 2). Performance improves with scale within the ESM2 family, with ESM2-650M achieving the best PCC (0.553), RMSE (7.430), and tail MAE (8.302), while ESM2-3B slightly improves MAE (4.920) but degrades on other metrics. Non-ESM2 models perform substantially worse. We therefore use ESM2-650M as the default backbone.

**Table 2.** Backbone comparison (test set).

| Model | PCC↑ | MAE↓ | RMSE↓ | Tail MAE↓ |
| --- | --- | --- | --- | --- |
| ESM2-8M | 0.336 | 5.825 | 8.550 | 9.706 |
| ESM2-35M | 0.443 | 5.491 | 8.049 | 9.191 |
| ESM2-150M | 0.505 | 5.265 | 7.750 | 8.617 |
| **ESM2-650M** | **0.553** | 4.964 | **7.430** | **8.302** |
| ESM2-3B | 0.540 | **4.920** | 7.542 | 8.664 |
| ProtT5 [1] | 0.417 | 5.366 | 8.137 | 9.688 |
| ESMC (300M) [2] | 0.399 | 5.690 | 8.261 | 9.227 |
| ESMC (600M) [2] | 0.414 | 5.768 | 8.268 | 9.517 |
| LC-PLM [16] | 0.175 | 6.213 | 9.389 | 12.049 |

#### Design choices

We ablate loss function, auxiliary supervision, and pooling (Table 3). The best regression performance is obtained with *l*_1_ loss without auxiliary supervision and with mean pooling, achieving the highest PCC (0.625) and lowest MAE (3.924) and RMSE (6.347). Auxiliary supervision does not improve standard regression metrics at this stage.

**Table 3.** Validation run ablations for ESM2-650M comparing loss (Huber vs. *l*_1_), auxiliary head (aux vs. no aux), and mean pooling (M) at auto-research epoch 17 (best performing configuration). Reported values are mean±std over 3 runs.

| Variant | PCC↑ | MAE↓ | RMSE↓ |
| --- | --- | --- | --- |
| Huber, no aux, M | 0.614±0.004 | 3.978±0.043 | 6.387±0.029 |
| Huber, aux, no M | 0.622±0.010 | 4.000±0.061 | 6.422±0.124 |
| Huber, aux, M | 0.616±0.009 | 3.966±0.013 | 6.371±0.044 |
| L1, no aux, M | **0.625±0.010** | **3.924±0.049** | **6.347±0.056** |
| L1, aux, M | 0.614±0.007 | 3.957±0.033 | 6.414±0.017 |

For tail performance (Table 4), the Huber + auxiliary + mean pooling variant achieves the lowest tail errors, indicating a trade-off between global regression accuracy and robustness on extreme examples.

**Table 4.** Tail error analysis for the ablation variants (validation set, auto-research epoch 17 (best performing configuration)). Tail MAE is the MAE on the top 10% largest |*ΔT*_*m*_| cases. Neg Tail MAE and Pos Tail MAE are the MAE on the most destabilizing and most stabilizing subsets, respectively. All variants use the ensemble head. M denotes mean pooling, and aux denotes the auxiliary classification head. Bold values indicate the lowest (best) error in each column. Reported values are mean±std over 3 runs.

| Variant | Tail MAE↓ | Neg Tail MAE↓ | Pos Tail MAE↓ |
| --- | --- | --- | --- |
| Huber, no aux, M | 7.473±0.094 | 7.175±0.106 | 10.477±0.143 |
| Huber, aux, no M | 7.639±0.257 | 7.375±0.286 | **10.291±0.040** |
| Huber, aux, M | **7.367±0.034** | **7.068±0.033** | 10.368±0.051 |
| L1, no aux, M | 7.416±0.088 | 7.120±0.100 | 10.388±0.189 |
| L1, aux, M | 7.503±0.015 | 7.213±0.025 | 10.420±0.116 |

#### Fine-tuning

We study the effect of fine-tuning depth (Table 5). Full finetuning yields the best performance. Partially unfreezing the top layers reduces training cost with moderate loss in accuracy, while freezing the backbone leads to substantial degradation.

**Table 5.** Fine-tuning regimes (test set).

| Regime | PCC↑ | MAE↓ | RMSE↓ | Tail↓ | Time(s) |
| --- | --- | --- | --- | --- | --- |
| **Full** | **0.553** | **4.964** | **7.430** | **8.302** | 2620 |
| Partial (8) | 0.516 | 5.148 | 7.681 | 8.543 | 1405 |
| Frozen | 0.467 | 5.486 | 8.037 | 8.515 | 1013 |

## 6 Discussion

Our results show that a carefully tuned sequence-only model can match or exceed more complex approaches for *ΔT*_*m*_ prediction on the S4346/S571 benchmark. The strong performance of ESM2-650M suggests that large protein language models encode sufficient information from sequence alone to support accurate prediction, without requiring explicit structural inputs.

### Ablations reveal key training choices

Our controlled ablation studies (Tables 3–4) highlight on the key contributors to the observed performance of our model. The best validation results are obtained with *l*_1_ loss, no auxiliary head, and mean pooling, making us believe that robust regression objectives and simple global aggregation are effective. While auxiliary supervision does not improve overall regression metrics, it can enhance sensitivity to extreme cases when combined with Huber loss. Also, mean pooling consistently improves performance, supporting the importance of global sequence context.

The fine-tuning study (Table 5) further highlights the importance of optimization strategy. Full fine-tuning performs best, whereas partial unfreezing retains reasonable performance at reduced cost, and freezing the backbone leads to a clear degradation. These findings indicate that effective adaptation of pretrained representations is critical for this task.

### Sequence homology drives the val–test gap

Validation performance is substantially better than test performance (MAE 3.92 vs. 4.96, see Table 1 and Table 3, respectively), which we attribute to differences in sequence homology. As described in Section 4.1, both training and validation sets are sampled from dataset `S4346` whereas the test set comes from dataset `S571`. As a consequence, we performed an in-depth analysis on both datasets by employing MMseqs2 `easy-search` (default sensitivity, E-value *<* 0.001). Here, we found that only 30% of the 37 test proteins have detectable homologs in the training set, while 70% represent novel protein families. In contrast, the validation split is derived from the same S4346 distribution, yielding a less severe distribution shift and, hence, to a better performance. This discrepancy largely explains the observed val–test gap and highlights the importance of evaluating generalization beyond closely related sequences.

### Comparison to prior work and metric trade-offs

Table 1 summarizes the comparison to prior methods. Our sequence-only model achieves the lowest MAE and RMSE among all approaches, despite not using pH or structural inputs, while remaining competitive in PCC.

DPStab [15] attains the highest PCC (0.592) but yields lower MAE and RMSE, illustrating a trade-off between ranking quality and absolute calibration. This difference can be traced to training and inference choices: MSE loss encourages regression toward the mean, compressing the prediction range and preserving relative ordering, while antisymmetric test-time ensembling further stabilizes rankings. In addition, DPStab leverages pH and implicit structural signals via ESM2 contact representations, which may contribute to the highest PCC correlation.

More broadly, Table 1 highlights a trade-off between correlation (PCC) and mean absolute error across methods. While some approaches that incorporate additional signals such as pH or structural information—most notably DP-Stab—achieve higher PCC, these gains do not consistently translate into improved MAE or RMSE.

In contrast, our approach uses full fine-tuning and *ℓ*_1_ loss, directly optimizing absolute error and avoiding mean-regression effects. This yields better calibrated predictions with lower MAE/RMSE at the cost of slightly reduced PCC. Overall, these results suggest a fundamental trade-off in *ΔT*_*m*_ prediction: objectives and strategies that favor smoothing and denoising improve ranking, whereas robust loss functions and end-to-end optimization improve calibration. Notably, our model achieves the best error metrics without additional input modalities, underscoring the importance of training design.

### Implementation of ESM2-650M in biopharma discovery workflows

By rational training of the state-of-the-art protein language model ESM2-650M with the S4346/S571 dataset to predict melting temperature changes upon mutation, we have achieved a functional model which can explore the mutation space and suggest successful mutations to improve the thermal stability of protein drug candidates, based on sequence input only. This is the case e.g., of early biopharmaceutical drug discovery phases, when only the sequence and few experimental measurements are known. Embedding of such models in discovery workflows at early stages, is a significant advancement in the drug discovery field, allowing prompt prioritization of designed mutants to decrease times and increase experimental throughput of workflows for large molecules.

## 7 Conclusion

We presented a study on *ΔT*_*m*_ prediction that challenges the need for structural inputs by showing that a carefully trained sequence-only model can achieve state-of-the-art results. Through an autoresearch-inspired setup search and a series of controlled ablations, we identified key factors that influence performance: using a large pre-trained backbone (ESM2-650M), applying mean pooling, choosing an appropriate loss function, and fully fine-tuning the model are all important contributors to strong accuracy. Our best sequence-based model achieved superior error metrics compared to published approaches that utilize protein structures or additional inputs, underscoring that *training strategy plays a crucial role* in this task. We hope that this work encourages a re-examination of how much value structural data adds for *ΔT*_*m*_ prediction, and highlights the considerable potential of sequence-based models when optimized for the task.

## Limitations

Several limitations remain. First, evaluation is restricted to a single benchmark split (S4346/S571), which limits statistical power and diversity. Second, we consider only single-point mutations, while practical protein engineering often involves combinatorial variants. Third, the sequence-only design is intentional; incorporating additional signals such as pH or structure could further improve performance, but is outside the scope of this study. Finally, antisymmetry-aware losses did not yield improvements in our experiments.

## References

1. Elnaggar, A., Heinzinger, M., Dallago, C., Rihawi, G., Wang, Y., Jones, L., Gibbs, T., Feher, T., Angerer, C., Steinegger, M., Bhowmik, D., Rost, B.: Prot-trans: Towards cracking the language of life’s code through self-supervised learning. IEEE Transactions on Pattern Analysis and Machine Intelligence (2021). 10.1109/TPAMI.2021.3095381

2. ESM Team: ESM Cambrian: Revealing the mysteries of proteins with unsupervised learning (2024), https://evolutionaryscale.ai/blog/esm-cambrian

3. Harmalkar, A., Rao, R., Xie, Y.R., Honer, J., Deisting, W., Anlahr, J., Hoenig, A., Czwikla, J., Sienz-Widmann, E., Rau, D., Rice, A.J., Riley, T.P., Li, D., Catterall, H.B., Tinberg, C.E., Gray, J.J., Wei, K.Y.: Toward generalizable prediction of antibody thermostability using machine learning on sequence and structure features. mAbs 15, 2163584 (2023). 10.1080/19420862.2022.2163584

4. Jia, L., Jain, M., Sun, Y.: Improving antibody thermostability based on statistical analysis of sequence and structural consensus data. Antibody Therapeutics 5, 202 (7 2022). 10.1093/ABT/TBAC017

5. Karpathy, A.: autoresearch: Llms doing autoresearch. https://github.com/karpathy/autoresearch (2026), accessed: 2026-03-27

6. Lin, Z., Akin, H., Rao, R., Hie, B., Rives, A., et al.: Evolutionary-scale prediction of atomic-level protein structure with a language model. Science 379(6637), 1123–1130 (2023). 10.1126/science.ade2574

7. Loshchilov, I., Hutter, F.: Decoupled weight decay regularization. In: International Conference on Learning Representations (2019)

8. Masso, M., Vaisman, I.I.: AUTO-MUTE: web-based tools for predicting stability changes in proteins due to single amino acid replacements. Protein Engineering, Design & Selection 23(8), 683–687 (2010). 10.1093/protein/gzq042

9. Moldwin, A.R., Shehu, A.: How much does protein structure really help? a case study in mutation-induced stability prediction. bioRxiv (2025). 10.64898/2025.12.22.694225, preprint

10. Natali, E., Hersch, J., Freiberg, C., Steigele, S.: Advancing large-molecule discovery with a unified digital platform for data analysis and workflow management. mAbs 17 (12 2025). 10.1080/19420862.2025.2555346

11. Pucci, F., Bourgeas, R., Rooman, M.: Predicting protein thermal stability changes upon point mutations using statistical potentials: Introducing HoTMuSiC. Scientific Reports 6, 23257 (2016). 10.1038/srep23257

12. Pudžiuvelytė, I., Olechnovič, K., Godliauskaitė, E., Sermokas, K., Urbaitis, T., Gasiunas, G., Kazlauskas, D.: Temstapro: protein thermostability prediction using sequence representations from protein language models. Bioinformatics 40(4), btae157 (2024). 10.1093/bioinformatics/btae157

13. Rives, A., Meier, J., Sercu, T., Goyal, S., Fergus, R., et al.: Biological structure and function emerge from scaling unsupervised learning to 250 million protein sequences. Proceedings of the National Academy of Sciences 118(15), e2016239118 (2021). 10.1073/pnas.2016239118

14. Steinegger, M., Söding, J.: Mmseqs2 enables sensitive protein sequence searching for the analysis of massive data sets. Nature Biotechnology 35(11), 1026–1028 (2017)

15. Wang, W., Zhou, Y., Huang, X., Wu, Y., Li, M., Zhang, Y.: DPStab: accurate prediction of protein stability changes from single mutations using self-distillation and antisymmetric constraint strategies. bioRxiv (2025). 10.1101/2025.05.18.654422, preprint

16. Wang, Y., Wang, Z., Sadeh, G., Zancato, L., Achille, A., Karypis, G., Rangwala, H.: LC-PLM: Long-context protein language modeling using bidirectional mamba with shared projection layers. Transactions on Machine Learning Research (TMLR) (2025)

17. Xu, Y., Liu, D., Gong, H.: Improving the prediction of protein stability changes upon mutations by geometric learning and a pre-training strategy. Nature Computational Science (2024). 10.1038/s43588-024-00716-2, published online 25 Oct 2024

18. Zhang, D., Zeng, Y., Hong, X., Xu, J.: Leveraging multi-modal representations to predict protein melting temperatures. In: Proceedings of the 39th AAAI Conference on Artificial Intelligence (Workshop on Foundation Models for Biology) (2025)

